# Myc inhibition triggers GM-CSF-driven regression of pancreatic tumours

**DOI:** 10.64898/2026.02.24.707647

**Authors:** T Campos, RM Kortlever, NM Sodir, MD Buck, J Stockis, J Parker, CM Lam, JR Patiño-Mercau, A Perfetto, A Edwards, S Boeing, TYF Halim, GI Evan

## Abstract

The complex aetiology of pancreatic ductal adenocarcinoma (PDAC), together with its desmoplastic and hypoxic microenvironment, hinders effective treatment and contributes to its rapid lethality^1^. Using a reversibly switchable genetic mouse model that closely recapitulates human PDAC phenotype progression, we previously showed that selective deactivation of oncogenic Myc in PDAC epithelial cells triggers rapid disassembly of advanced PDACs^2^, both tumor cells and their associated immune microenvironment. Here, leveraging spatiotemporal genomics and multiplex immune profiling we determine the mechanism underpinning this regression programme and identify granulocyte-macrophage colony-stimulating factor (GM-CSF) as its key instructive cytokine, transiently released by pancreatic ductal epithelial cells rapidly following Myc inactivation, that initiates tumour regression. We further demonstrate that type 1 conventional dendritic cells (cDC1s) act as critical effectors in Myc-OFF GM-CSF-driven tumour regression. Both antibody-mediated blockade of GM-CSF and genetic ablation of cDC1s via *Batf3* knockout bone marrow transplantation prevent PDAC regression. Conversely, transient systemic administration of recombinant GM-CSF to PDAC-bearing mice promotes rapid cDC1 infiltration and induces marked tumour regression in the sustained presence of Myc activity. Together, these findings reveal that PDAC regression induced by Myc de-activation is mediated by a latent morphogenic programme that is initiated by transient release of GM-CSF from tumour cells. This regression programme is rapid, tissue-specific, involves both neoplastic cells and attendant desmoplastic stroma, is reliant on innate immunity and provides a novel framework for therapeutic intervention in PDAC.

## INTRODUCTION

Pancreatic ductal adenocarcinoma (PDAC) remains one of the deadliest human cancers, with a 5-year survival rate below 9% and little improvement in the past 3 decades^3^. Its grim prognosis stems largely from its late clinical presentation as an already invasive disease together with its profound innate resistance to conventional chemotherapy and radiotherapy. PDAC is characterized by a dense fibroinflammatory desmoplasia that arises through complex interactions between tumour cells and adjacent mesenchymal, endothelial, inflammatory, immune, and neuronal cells^1,4^. This dense stroma is thought to contribute to PDAC’s therapeutic resistance by restricting vascular perfusion and oxygenation^5,6^ and suppressing antitumour immune responses^7^. Pathological analyses of resected tumours indicate that overt PDAC arises from pancreatic intraepithelial neoplasms (PanINs), indolent precursor lesions classified into four stages (PanIN-1A, -1B, -2, and -3) based on increasing nuclear and architectural atypia^8–10^. At the genetic level, PDAC is associated with a signature ensemble of recurring oncogenic mutations. Most notably, oncogenic KRas mutations are present in over 90% of PanINs and are generally thought to be founder events^11^ but even those relatively few PDACs that carry no KRas mutations show evidence of oncogenic mutations in the wider “Ras pathway”^12^. The prototypical cooperating oncogenic partner of KRas is Myc^13^, a pleiotropic transcription factor whose putative role is to coordinate expression of the thousands of genes that, together, coordinate the diverse genetic programmes needed for cell proliferation and is tightly regulated in normal somatic cells by mitogen availability^2^. In contrast, aberrantly persistent, deregulated Myc is implicated in the majority, perhaps all, cancers. Such deregulation is usually an indirect consequence of Myc’s relentless induction by “upstream” oncogenes or mutations in regulatory enhancers within the *MYC* gene^14–20^. In high grade cancers, including PDAC, the *MYC* gene is also sometimes amplified ^21^. Diverse studies indicate a critical role for Myc in both the progression from PanIN to PDAC and in subsequent PDAC maintenance^5,16,22–24^. Accordingly, we previously reported in a reversibly switchable genetic mouse model of PDAC, driven by oncogene cooperation of KRas*^G12D^*and Myc, that acute activation of Myc triggers immediate and synchronous progression of pancreatic tumours that, from the outset, exhibit the signature characteristics of PDAC — including rapid growth, prolific desmoplasia, hypoxia and microvascular occlusion, local invasiveness, and in a range of morphological PDAC subtypes (e.g. classical, mucinous, spindle-cell, mixed phenotype)^2^. The switchability afforded by our mouse model indicated that all these features are completely reversible upon de-activation of Myc, with a notably rapid in the immune compartment of the microenvironment that preceded overt tumour regression. Here, we define the nature of the PDAC regression programme, identifying the key players that initiate and instruct tumour cell death and deconstruction of desmoplastic tissue, and thereby uncovering novel potential therapeutic targets for treating pancreatic cancer.

## RESULTS

### Determining rapidity of pancreatic adenocarcinoma regression onset after Myc OFF

To investigate in detail both the progression and regression of PDAC *in vivo* we used our previously described *p48-Cre; LSL-KRas^G12D^; Rosa26LSL-MycER^T^*^2^ (*KM*) mouse model, in which oncogenic KRas^G12D^ is expressed in early pancreas ontogeny by the “hit-and-run” action of Cre recombinase driven from the *Ptf1/p48* promotor. The same Cre switch simultaneously induces expression of the reversibly activatable, 4-hydroxyTamoxifen (4-OHT)-dependent allele of Myc — MycER*^T^*^2^ ^25^. Notably, KRas^G12D^ and MycER*^T^*^2^ (hereafter Myc) are both expressed at quasi-physiological levels^26^. As previously reported, in the absence of Myc activity *KM* mice develop multiple PanIN foci that almost all remain as indolent lesions for the host’s lifetime. In contrast, acute and sustained activation of ectopic Myc via systemic administration of tamoxifen (TAM) in the diet^27^, as depicted in Fig. 1a, triggers rapid transition of PanINs into highly proliferative and aggressive pancreatic adenocarcinomas (PDAC) that share all the signature phenotypic features of human PDAC — including pervasive desmoplasia, vascular stenosis and hypoxia, inflammation, immune suppression and invasiveness^2^.

**Figure 1.**
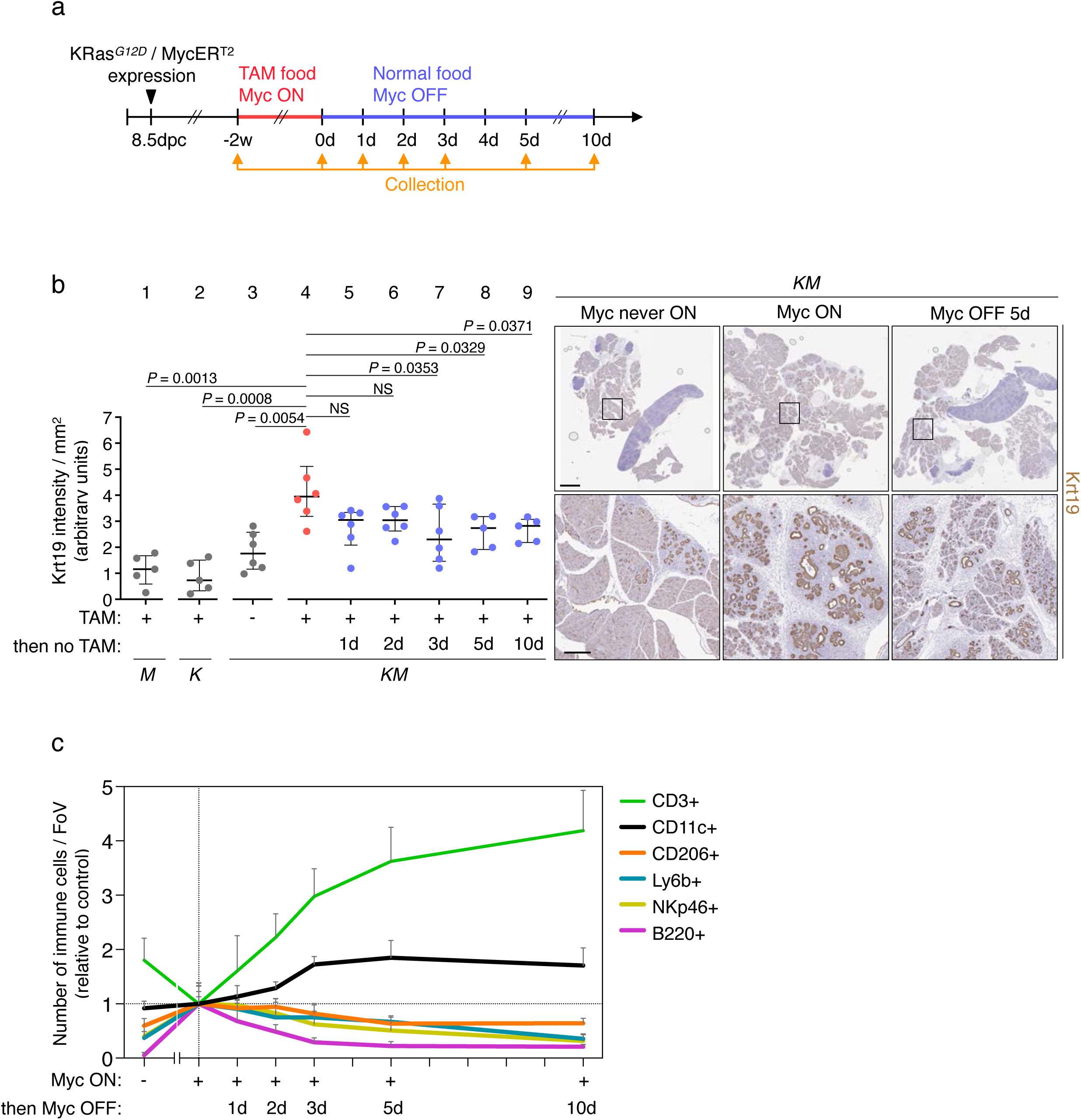
Oncogenic Myc de-activation triggers immediate onset of pancreatic adenocarcinoma regression. **a.** Experimental design: 12-weeks-old *KM* mice were treated with tamoxifen food for 2 weeks to activate MycER*^T^*^2^, followed by treatment with normal food to deactivate MycER*^T^*^2^. Samples were collected at the indicated time points. dpc= days postcoitum, w= weeks, d= days. **b.** Left: Keratin 19 (Krt19) immunohistochemistry staining quantification of pancreata from *M, K* and *KM* mice treated with tamoxifen food (TAM) for 2 weeks (+), normal food (-) or tamoxifen food for 2 weeks followed by normal food (then no TAM) for the indicated time points. N = 5-6 mice per group. Shown are medians with interquartile ranges per group. Statistical analysis was performed using multiple unpaired *t*-tests with Welch’s correction. Right: representative Krt19 staining of pancreata from *KM* mice with normal food (Myc never ON), tamoxifen food for 2 weeks (Myc ON), or tamoxifen food for 2 weeks followed by normal food for 5 days (Myc OFF 5d). Scale bar across the top row = 2 mm. Bottom row shows higher magnification of the insets in top row, scale bar across the row = 300 μm. **c.** Superimposed quantification of CD3+, CD11c+, CD206+, Ly6b+, NKp46+ and B220+ cells immunostainings in pancreata from *KM* mice treated with normal food (-), tamoxifen food (Myc ON) for 2 weeks (+) or tamoxifen food for 2 weeks followed by normal food (then Myc OFF) for the indicated time points. FoV = field of view. N = 5-8 mice per timepoint. Shown are means and standard deviations per group.

To delineate the extent of tumour growth we used cytokeratin 19 (Krt19) as immunohistochemical marker for pancreatic tumour epithelium. Although Krt19 is normally expressed in the lining of the gastroenteropancreatic and hepatobiliary tracts, it also serves as a dynamic marker for early and late stage PDAC^28^. Pancreas sections from mice in which Myc had been persistently activated for 2 weeks showed significant increase in Krt19 positive cells (Fig. 1b, 4^th^ column in graph, *KM* mice treated with TAM) relative to control samples (Fig. 1b, 1^st^ to 3^rd^ column in graph, *M* and *K* mice treated and *KM* not treated with TAM). Detailed analysis of the PDACs microenvironment at 2 weeks of Myc ON revealed the tumours to be infiltrated by diverse immune cells, including anti-inflammatory macrophages (CD206+), neutrophils (Ly6b+), NK cells (NKp46+) and B cells (B220+) (Fig. 1c and 2). Myc was then de-activated in pancreata of PDAC-bearing mice by withdrawal of tamoxifen. This induced rapid regression of PDAC from invasive tumour back to PanIN together with loss of Krt19+ tumour mass, evident by 1 day and significant by 3 days Myc OFF (Fig. 1b, 5^th^ to 9^th^ columns compared to 4^th^ column in graph), rapid proliferative arrest, as evidenced by loss of Ki67+ signal together with a rapid increase in tumour cell death (TUNEL+ staining), that peaked around day 3 to 5 after Myc OFF (Extended data Fig. 1a), and dramatic reorganization of the local PDAC tumour microenvironment: rapid exclusion of macrophages, neutrophils, NK cells and B cells from the tumour masses together with influx of T cells (CD3+ cells) in the vicinities of the tumour masses (Fig. 1c). We also observed local accumulation of CD11c+ cells (primarily myeloid derived cells) in the regions of the regressing tumours (Fig. 1c, black line and Extended data Fig. 1b). Somewhat delayed compared with the rapid changes to the immune and tumour cells death that Myc deactivation induces we observed shrinkage of the abundant desmoplasia that made up the bulk of PDAC tumour mass, as evident from H&E staining (Extended data Fig. 1c and 2). Our data indicate that the Myc-OFF–initiated morphogenic programme that orchestrates PDAC regression is multifaceted and involves many cell types, signals and interactions. Nonetheless, it Myc-OFF switch originates within the PDAC tumour cells themselves and it must therefore be transduced by some means to downstream effector cells that run the downstream regression programme.

**Figure 2.**
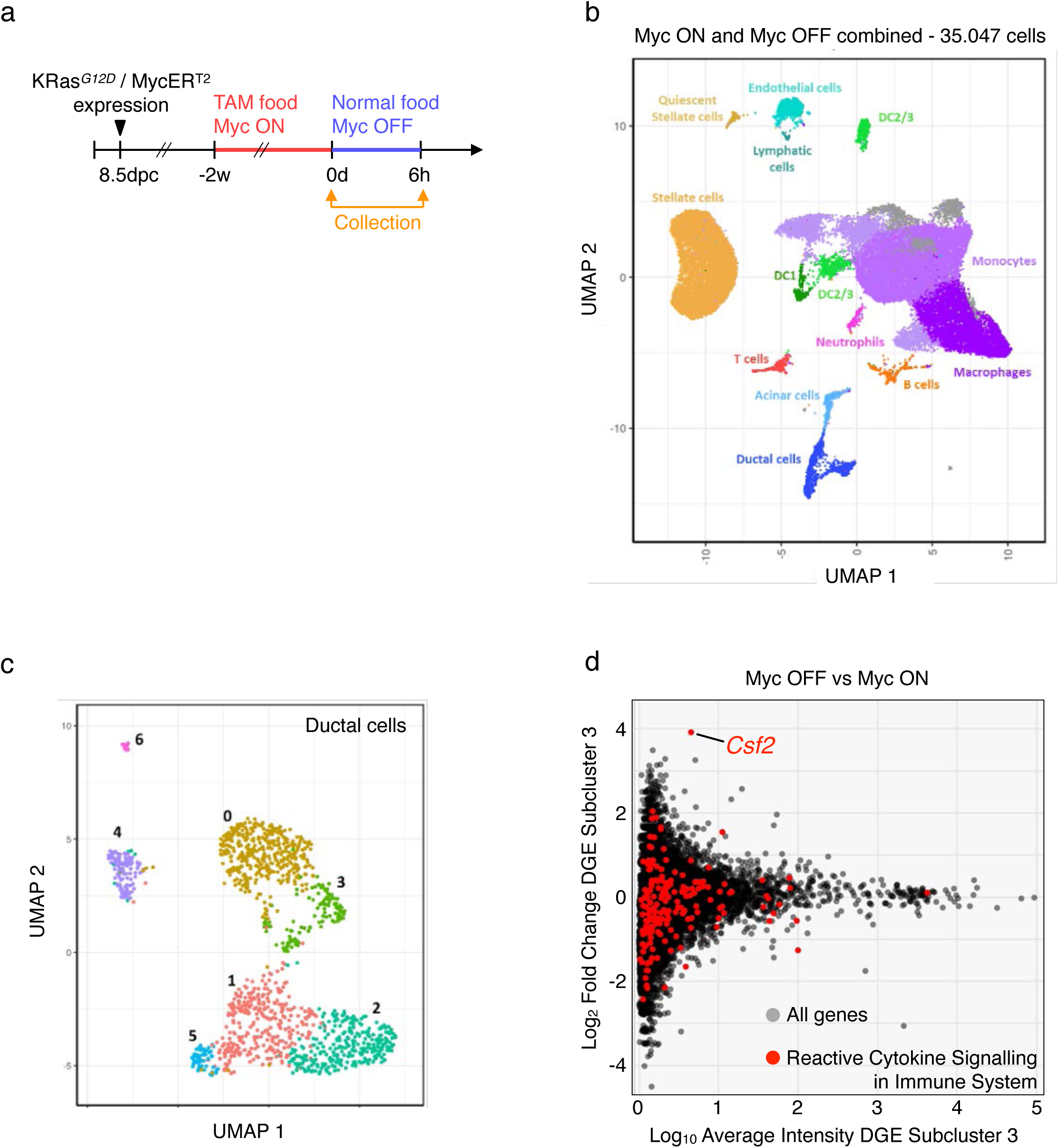
Acute loss of oncogenic Myc triggers rapid expression of *Csf2* in ductal tumour cells. **a.** Experimental design: 12-weeks-old *KM* mice were treated with tamoxifen food for 2 weeks to activate MycER*^T^*^2^ followed by normal food for 6 hours to deactivate MycER*^T^*^2^. Samples were collected at the indicated time points. **b.** Uniform manifold approximation and projection (UMAP) visualisation of single-cell RNA-sequence analysis plotting whole pancreas from *KM* mice treated with tamoxifen food for 2 weeks (Myc ON) and tamoxifen food for 2 weeks followed by normal food for 6 hours (Myc OFF). Cells are coloured and named by subpopulations. N = 2-3 mice per time point. **c.** UMAP visualization of single-cell RNA-sequence data plotting only the pancreatic ductal epithelial cell compartment (Krt19+ cells). Cells coloured by subpopulation clusters. **d.** Differential gene expression (DGE) analysis shown as MA plot (Log_2_ Fold Change versus Log_10_ Average Intensity) of ductal cells Subcluster 3 (related to Fig. 2c) comparing pancreata of mice treated with tamoxifen food for 2 weeks followed by normal food for 6 hours with pancreata of mice treated with tamoxifen food for 2 weeks. In red: Reactome Cytokine Signalling in Immune System genes (R-HSA-1280215). *Csf2* is highlighted.

To explore this initial signal relay, we first established how rapidly acute withdrawal of tamoxifen triggers a detectable fall in Myc transcriptional activity in *KM* pancreatic cells. To do this we first identified suitably short-lived Myc transcriptional targets that might serve as a readout of loss of Myc activity using bulk RNA-sequencing (RNA-seq) data previously obtained using a *KM* mouse-derived PDAC cell line (OmicsDI accession number E-MTAB-8467^2^). As described^2^, MycER*^T^*^2^ was activated and maintained in cultured *KM* cells by addition of 4-OHT to the growth medium. 4-OHT was then withdrawn to de-activate MycER*^T^*^2^ and at various time points thereafter RNA-seq was performed. We identified the most rapidly attenuated Myc target mRNA as *Rasd2* (RASD Family Member 2) (Extended data Fig. 2a). By analysing Myc ChIP-seq data from the same *KM* cell line (OmicsDI accession number GSE234078^29^), we confirmed that MycER binds the *Rasd2* promoter in the presence of 4-OHT, and that this interaction is markedly reduced by 24 hours post 4-OHT withdrawal (Extended data Fig. 2b). The short half-life of *Rasd2* was confirmed by *in situ* hybridisation analysis of pancreata from *KM* mice, and showed a rapid drop in signal by 12 hours post Myc OFF (Extended data Fig. 2c). Further qRT-PCR analysis confirmed the *in vivo* half-life of *Rasd2* RNA to be around 3 hours (Extended data Fig. 2d, vertical dotted line). Consequently, we used *Rasd2* as a biomarker for loss of Myc activity *in vitro* and *in vivo*. Given our previous studies showing that a 50% drop in physiological Myc level is sufficient to disengage Myc’s oncogenic activity^,26^, we decided on 6 hrs post Myc OFF as our time point to search for early instructive tumour regression signals.

### Myc de-activation triggers rapid release of *Csf2* in pancreatic epithelial cells

We next sought to identify candidate regression-initiating signals released from PDAC tumour cells in response to Myc OFF. First, to maintain MycER activity and drive formation of multiple PDACs, adult *KM* mice were treated continuously for 2 weeks with tamoxifen. Animals were then transferred to tamoxifen-free food for 6 hours (Myc OFF), as described in Fig. 2a, and single-cell RNA-seq performed to deconvolute gene expression in the mouse pancreata. Grouping all samples together, discrete cell subpopulations were mapped to specific cell types (Fig. 2b, Extended Data Fig. 3a and Extended Data Table 1 for full list of markers). Since MycER is exclusively expressed in epithelial cells, we then focused our analysis on cells from the epithelial compartment in which Myc was toggled OFF (Extended Data Fig. 3b) by subclustering the Krt19+ epithelial compartment (Fig. 2c and subclusters markers in Extended Data Fig. 3c). Differential gene expression analysis comparing 6 hours Myc OFF with Myc ON revealed *Csf2* (Colony-Stimulating Factor 2) to be the only immune-related signaling gene whose expression was clearly elevated after switching Myc OFF (Fig. 2d). Immunohistochemistry confirmed that GM-CSF (encoded by the *Csf2* gene) was induced in pancreatic epithelial cells at 6 hrs post Myc OFF (Extended Data Fig. 3d). Cytokine array analysis showed that the Myc OFF-induced increase in GM-CSF protein is transient and peaks at 6-12 hours after deregulated Myc downregulation. (Extended Data Fig. 3e). Together, these data validated GM-CSF’s candidacy as an initial instructive signal that engaged the regression pathway responsible for the Myc-OFF dependent-pancreatic tumour regression.

### GM-CSF is required for Myc blockade-induced regression of pancreatic tumours

GM-CSF is known to be critical in the development and function of myeloid-derived immune cells, including dendritic cells and macrophages^30,31^. To ascertain whether GM-CSF also plays an early causal role in pancreatic tumour regression after Myc OFF, we blocked its activity systemically using a GM-CSF-specific blocking antibody. Pancreatic adenocarcinomas were generated in *KM* mice by activating Myc for 2 weeks. Starting two days before switching Myc OFF, *KM* mice received either an *i.p.* bolus of anti-GM-CSF antibody or control matched IgG every other day until 10 days post-Myc OFF. PDAC samples were collected 3 and 10 days after switching Myc OFF (Fig. 3a). GM-CSF blockade profoundly impaired the regression response to Myc OFF. Compared with controls at 3 days Myc OFF (Fig. 3b, 2^nd^ and 4^th^ columns in graphs), animals treated with anti-GM-CSF antibody (Fig. 3b, 3^rd^ and 5^th^ columns in graphs) showed a greatly impaired tumour regression, evidenced by absence of fall in Krt19+ cells and of overall tumour burden (Fig. 3b, 1^st^ and 2^nd^ row). GM-CSF blockade also abrogated the transient wave of cell death (TUNEL+) previously observed 3 days after switching Myc OFF (Fig. 3b, 3^rd^ row). The impact of anti-GM-CSF on Myc-OFF-driven tumoural cell death was confirmed by quantification of double immunoassayed tissue for cleaved caspase 3 (CC3 - apoptosis marker) and Pan-Keratin (PanK - epithelial marker), showing a significant decrease in the number of epithelial cells dying (CC3+/PanK+) in pancreatic tumours of mice treated with anti-GM-CSF antibody relative to IgG-treated controls (Fig. 3c). The number of CD206+ macrophages, Ly6b+ neutrophils, NKp46+ NK cells, B220+ B cells or CD3+ T cells in the PDAC stroma was not always significantly or consistently impacted by GM-CSF blockade (Extended Data Fig. 4a-e). In contrast, whereas Myc OFF induced a profound influx of CD11c+ myeloid cells into pancreatic tumours, that influx was completely suppressed by anti-GM-CSF blockade (Fig. 3d, see also Fig. 1c and Extended Data Fig. 1c). Blocking GM-CSF had no detectable inhibitory impact on the profound proliferative arrest that deactivating Myc triggers in pancreas tumours, consistent with the notion that proliferation is privately regulated by Myc activity (Fig. 3b, 4^th^ row).

**Figure 3.**
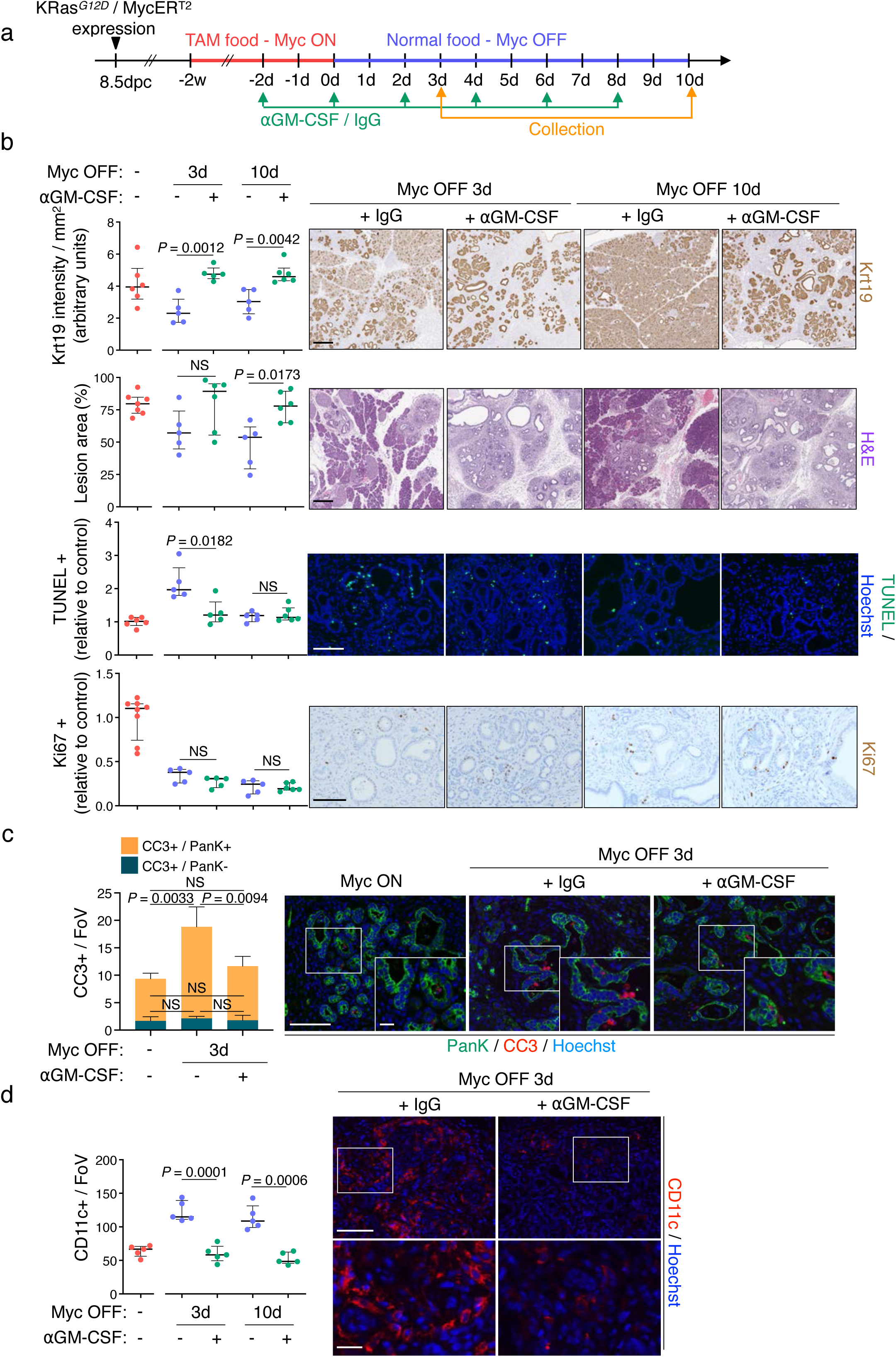
GM-CSF is required for PDAC regression. **a.** Experimental design: 12-weeks-old *KM* mice were treated with tamoxifen food for 2 weeks to activate MycER*^T^*^2^ followed by normal food to deactivate MycER*^T^*^2^. Blocking antibody for granulocyte-macrophage colony-stimulating factor (αGM-CSF) or IgG control was administered every other day as indicated. Samples were collected at the indicated time points. **b.** Left: Immunohistochemistry, haematoxylin and eosin or immunofluorescence quantification of pancreata from *KM* mice treated with tamoxifen food for 2 weeks (1^st^ column, data taken from Fig. 1b and Extended Data Fig. 1a and b) or tamoxifen food for 2 weeks followed by normal food for 3 or 10 days while treated with IgG control (2^nd^ and 4^th^ column) or αGM-CSF antibody (3^rd^ and 5^th^ column). Tissue sections stained for Keratin 19 (Krt19; 1^st^ row), haematoxylin and eosin (H&E; 2^nd^ row), Cell death (TUNEL; 3^rd^ row) and proliferation (Ki67; 4^th^ row). N = 5-6 mice per group. Right: representative staining as indicated at the top of panels. Scale bars across the row = 300 μm for Krt19 and H&E, 100 μm for TUNEL and Ki67. **c.** Left: Pan-keratin (PanK; green) and cleaved caspase 3 (CC3; red) immunofluorescence staining quantification of pancreata harvested from *KM* mice treated with tamoxifen food for 2 weeks or tamoxifen food for 2 weeks followed by normal food for 3 days while treated with IgG control or αGM-CSF antibody. N = 5-6 mice per group. Right: representative immunostaining for indicated conditions / timepoints, scale bar across the row = 100 μm. Insets show higher magnification of a section of the corresponding panel, scale bar across the row = 20 μm. **d.** Left: CD11c immunofluorescence staining quantification of pancreata harvested from *KM* mice treated with tamoxifen food for 2 weeks (1^st^ column, data taken from Fig. 1c) or tamoxifen food for 2 weeks followed by normal food for 3 or 10 days while treated with IgG control (2^nd^ and 4^th^ column) or αGM-CSF antibody (3^rd^ and 5^th^ column). N = 5 mice per group. Right: representative immunostaining for indicated conditions at 3 days Myc OFF. Scale bar across the row = 100 μm. Bottom row shows higher magnification of the corresponding insets in top panels, scale bar across the row = 20 μm. **b, c, d:** Shown are median with interquartile range per group. Statistical analysis was performed using multiple unpaired *t*-tests with Welch’s correction (Krt19, TUNEL, Ki67, CC3, CD11c) or multiple Mann-Whitney tests (H&E).

### Exogenously administered GM-CSF is sufficient to trigger the PDAC regression programme

Having established that GM-CSF is necessary downstream of the Myc OFF-induced PDAC regression programme, we next asked whether it is sufficient. *KM* mice were kept on tamoxifen diet for 17 days to maintain Myc active and generate multifocal pancreatic adenocarcinomas. On days 13 and 14, tumour-laden animals were intraperitoneally injected with either recombinant GM-CSF (rGM-CSF) or vehicle control, emulating the timing of the GM-CSF peak as observed during Myc OFF regression, pancreas tissue was collected 3 days later (Fig. 4a). Compared with control *KM* mice, animals treated with rGM-CSF showed profound reduction in Krt19 staining and tumour burden (Fig. 4b, 1^st^ and 2^nd^ row), as well as substantially elevated cell death (Fig. 4b, 3^rd^ row). We then confirmed that the cells dying due to the administration of rGM-CSF were predominantly epithelial (Fig. 4c). Ectopic rGM-CSF administration elicited no change in overall cell proliferation (Fig. 4b, 4^th^ row), confirming that Myc controls proliferative status of the regressing tumour independently of the GM-CSF-driven regression programme. Further analysis of the pancreatic tumoural area indicated a substantial increase in both CD11c+ and CD3+ cells in mice treated with rGM-CSF (Fig. 4d and Extended Data Fig. 5a, respectively), mirroring the dramatic influx of CD11c+ and CD3+ T cells originally observed upon Myc deactivation in PDACs (Fig. 1c). Administration of rGM-CSF did not significantly affect the distribution of other microenvironmental immune cells analysed (Extended Data Fig. 5b-e). In summary, our data indicate that, despite sustained Myc activity, exogenous rGM-CSF is sufficient to drive key phenotypic attributes of PDAC regression.

**Figure 4.**
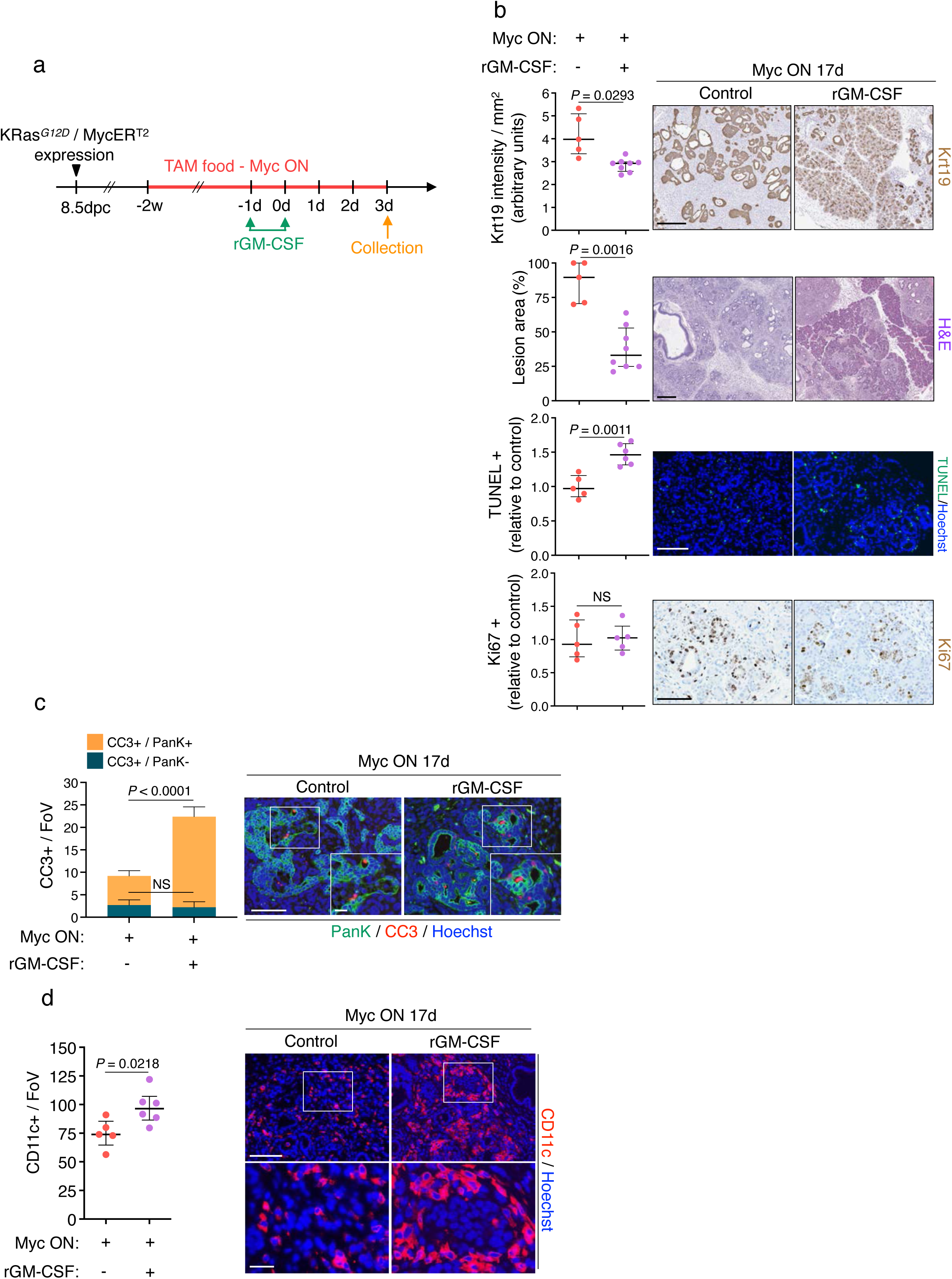
Systemic recombinant GM-CSF treatment is sufficient to enforce PDAC regression. **a.** Experimental design: 12-weeks-old *KM* mice were treated with tamoxifen food to activate MycER*^T^*^2^. Recombinant granulocyte-macrophage colony-stimulating factor (rGM-CSF) or vehicle control was administrated on days 13 and 14. Samples were collected 3 days later (Myc ON 17 days). **b.** Left, graphs: Immunohistochemistry and haematoxylin and eosin (H&E) quantification of pancreata from *KM* mice treated with tamoxifen food for 17 days (1^st^ column) or tamoxifen food for 17 days and treated with rGM-CSF (2^nd^ column). Tissue sections stained for Keratin 19 (Krt19; 1^st^ row), H&E (2^nd^ row), cell death (TUNEL; 3^rd^ row) and proliferation (Ki67; 4^th^ row). N = 5-8 mice per group. Right: representative sections staining. Scale bars across the row = 300 μm for Krt19 and H&E, 100 μm for TUNEL and Ki67. **c.** Left: Pan-keratin (PanK; green) and cleaved caspase 3 (CC3; red) immunofluorescence quantification of pancreata harvested from *KM* mice treated with tamoxifen food for 17 days (1^st^ column) or tamoxifen food for 17 days and treated with rGM-CSF (2^nd^ column). N = 5-8 mice per group. Right: representative immunostaining for indicated conditions / timepoints, scale bar across the row = 100 μm. Insets show higher magnification of a section of the corresponding panel, scale bar across the row = 20 μm. **d.** Left: CD11c immunofluorescence staining quantification of pancreata harvested from *KM* mice treated with tamoxifen food for 17 days (1^st^ column) or tamoxifen food for 17 days and treated with rGM-CSF (2^nd^ column). N = 5-6 mice per group. FoV = field of view. Right: representative sections immunofluorescence staining. Scale bar across the row = 100 μm. Bottom row shows higher magnification of the corresponding insets in top panels, scale bar across the row = 20 μm. **b, c, d:** Shown are median with interquartile range per group. Statistical analysis was performed using multiple unpaired *t*-tests with Welch’s correction (Krt19, TUNEL, Ki67, CC3, CD11c) or multiple Mann-Whitney tests (H&E).

### Type 1 dendritic cells are required for Myc OFF-induced PDAC regression

We established that Myc downregulation triggers PDAC tumour cells to transiently release GM-CSF, which then presumably signals to CD11c+; GM-CSFR+ target cells located in the stromal milieu of the pancreatic tumour. This, in turn, then engages the downstream morphogenic cascade of tumour regression. Multiple immune cell types weakly or transiently express the CD11c marker, including T cells, NK cells, monocytes, macrophages, neutrophils and B cells^32^. By contrast, CD11c is consistently expressed at high levels by conventional dendritic cells (cDCs), especially type 1 (cDC1)^32,33^. Flow cytometric analysis confirmed a significant relative increase in cDC1 cells over total of dendritic cells following Myc-OFF (Extended Data Fig. 6a). To validate cDC1s as the immediate downstream targets of GM-CSF in Myc OFF induced PDAC regression programme, we used CD45.1;*Batf3-/-* mice, which lack BATF3, the essential transcription factor for the development of the cDC1 lineage^34^. To construct *KM* mice with bone marrow unable to generate cDC1s, we generated a radiation bone marrow chimeras (Fig. 5a): *KM* mice expressing the variant CD45.2 allele (CD45.2;*KM*) were irradiated to ablate their bone marrow and then used as recipients for bone marrow from CD45.1;*Batf3-/-* mouse donor (resulting in *KM:Batf3-/-* mice). As a donor control, we used CD45.1;*WT* mouse (*KM:WT*). 5 weeks after bone marrow reconstitution both *KM:Batf3-/-* and *KM*:*WT* chimeric mice displayed more than 90% CD45.1+ donor immune cells (Extended Data Fig. 6b, left graph). As expected, bone marrow reconstituted *KM:Batf3-/-* mice exhibited sharply reduced numbers of cDC1 cells (0.015% of total CD45+ cells) compared to a 10-fold higher proportion of cDC1 cells in *KM:WT* control mice (0.111% of total CD45+ cells) (Extended Data Fig. 6b, right graph). To ascertain whether cDC1 cells play a key role in PDAC regression, reconstituted *KM:Batf3^-/-^* mice were treated with tamoxifen food for 2 weeks (Myc ON) and then shifted to normal food for 3 days (Myc OFF) (Fig. 5a). Compared with Myc ON (Fig. 5b, left columns), mice in which Myc was switched OFF for 3 days (Fig. 5b, right columns in graphs) showed no discernible reduction in Krt19+ PDAC tumour cells, no reduction in overall tumour burden and no increase in TUNEL+ cell death (Fig. 5b, 1^st^, 2^nd^ and 3^rd^ row), indicating that pancreatic tumour regression was greatly impaired in *KM* mice lacking cDC1+ cells. Nonetheless, we still observed the expected substantial reduction in tumour cell proliferation due to the direct influence of oncogenic Myc shutdown (Fig. 5b, 4^th^ row). In contrast, control *KM:WT* chimeric mice showed a significant increase in cell death after switching Myc OFF (Extended Data Fig. 6c), indicating that the PDAC regression programme was still ongoing and had not been perturbed by the bone marrow transplant procedure itself. Thus, we conclude that cDC1 cells are required for the Myc-OFF-dependent regression of pancreatic tumours.

**Figure 5.**
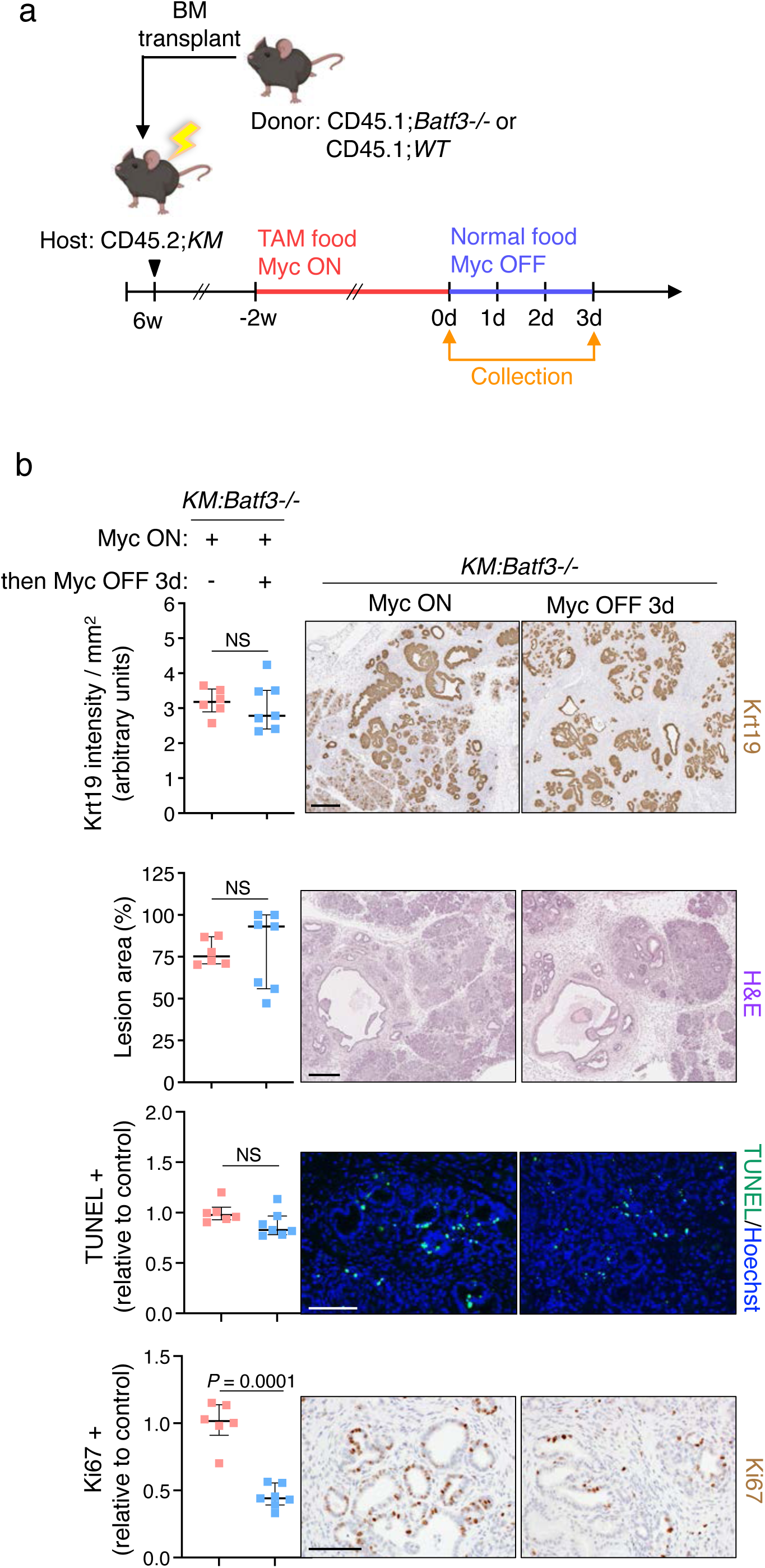
cDC1s are required for Myc OFF-induced regression of pancreatic tumour cells. **a.** Experimental design: 6-weeks-old recipient *KM* mice were irradiated and bone marrow from donor mice transplanted the following day (resulting in chimeric *KM:Batf3^-/-^* or *KM:WT* mice). 14-weeks-old chimeric mice were treated with tamoxifen food for 2 weeks to activate MycER*^T^*^2^ followed by normal food to deactivate MycER*^T^*^2^. Samples were collected at indicated time points. **b.** Left: immunostaining quantification of pancreata from *KM:Batf3-/-* mice treated with tamoxifen food for 2 weeks (Myc ON; 1^st^ column) or tamoxifen food for 2 weeks and then normal food for 3 days (2^nd^ column). Tissue sections stained for Keratin 19 (Krt19; 1^st^ row), haematoxylin and eosin (H&E; 2^nd^ row), cell death (TUNEL; 3^rd^ row) and proliferation (Ki67; 4^th^ row). N = 6-7 mice per group. Shown is median with interquartile range per condition/timepoint. Statistical analysis was performed using multiple unpaired *t*-tests with Welch’s correction (Krt19, TUNEL, Ki67) or multiple Mann-Whitney tests (H&E). Right: representative sections staining of Myc ON and Myc OFF 3d in *KM:Batf3-/-* mice. Scale bars across the row = 300 μm for Krt19 and H&E, 100 μm for TUNEL and Ki67.

## DISCUSSION

Previously, we sowed that withdrawal or blockade of oncogenic Myc activity in KRas*^G12D^/Myc* - dependent PDAC model triggers rapid tumour regression^2^. As confirmed in the present work, PDAC regression involves dynamic changes across all cell types within the tumour mass — epithelial, endothelial, mesenchymal, inflammatory and immune, and is characterised by a sharp fall in tumour cell proliferation, onset of tumour cell apoptosis, and progressive normalisation of pancreas glandular structure. We investigate why inhibiting the oncoprotein Myc — a universal driver of cell growth — triggers tumour regression and cell death in PDAC cancer cells, rather than merely pausing cancer cell proliferation.

In our PDAC model Myc is shut down only in the tumour cell compartment. We therefore reasoned that Myc blockade must trigger release of transcellular signals that originate in the MycER*^T^*^2^-positive tumour cells and then cascades across the pancreatic tumour milieu and instruct the process of tumour regression. The rapidity and tightly choreographed dismantling of the PDAC tumours that accompanies PDAC regression, together with the appearance of new characteristics, such as the sustained influx of CD11c+ cells, indicates that Myc-OFF induced PDAC regression is not merely a passive reversal of Myc-driven tumourigenesis but an active morphogenic programme in its own right. Using differential gene expression analysis, we uncovered a PDAC epithelial sub-cluster of cells that by 6 hrs post Myc OFF broadcasts a transient wave of the signalling molecule GM-CSF, the product of the gene *Csf2.* GM-CSF is a broadly active cytokine commonly associated with haematopoiesis, in particular the growth and development of granulocytes and macrophages. In response to inflammatory stimuli and cytokines it is expressed by a variety of cell types, including T cells, macrophages, endothelial cells, epithelial cells, and fibroblasts and plays important role in maintenance of integrity and damage repair in many tissues^35–41^. We established a causal role for GM-CSF in driving PDAC regression by demonstrating both its necessity and sufficiency. To confirm necessity, we showed that systemic blockade of GM-CSF suppresses the rapid Myc OFF-induced loss of PDAC tumour mass and abrogates the afore mentioned rapid and sustained influx of CD11c+ cells into the tumour masses.

Conversely, systemic administration of tumour-bearing *KM* mice to recombinant GM-CSF was sufficient to trigger major features of the PDAC regression programme, such as rapid loss of tumour cells and tumour burden and influx of both CD3+ and CD11c+ cells, even when Myc activity was maintained throughout. However, tumour cell proliferation remained, as ever, dependent solely on Myc activity, demonstrating once again uncoupling of proliferation control (Myc-dependent) from cell death (GM-CSF-dependent) in regression. Administration of either GM-CSF blocking antibody or recombinant protein to Kras^G12D^/Myc-driven PDAC bearing mice had no significant impact on the dynamics of other major haematopoietic cell types (NK, B220, neutrophil and CD206 macrophages). Our observation that exogenous administration of recombinant GM-CSF is sufficient to trigger PDAC regression phenotype raises the potentially therapeutic relevant implication that once oncogene inhibition has triggered GM-CSF release in at least some of the PDAC tumour cells, the regression programme can propagate to bystander tumour cells.

Having established GM-CSF as an output of the Myc OFF regression signalosome, we next searched for candidate sentinel cells that might act as a receiver and propagator of the GM-CSF-dependent regression signal. GM-CSFR is widely expressed by many cell lineages, both hematopoietic (e.g. neutrophils, eosinophils, monocytes and macrophages, dendritic cells and CD34^+^ progenitors) and non-hematopoietic (endothelial, epithelial, mesenchymal) cells. Almost all analysed immune cell types were spatially unperturbed by GM-CSF blockade or administration, except for CD3^+^ and CD11c+ cells. Because of their well attested roles in suppression of neoplasia, T cells were the most intuitively plausible cell to relay the immediate GM-CSF regression programme. However, we had previously used systemic antibody treatment to ablate both CD4^+^ and CD8^+^ T cells and shown that this did not measurably inhibit PDAC regression^2^. It nonetheless remains possible that some rare type of CD3-positive CD4^-^ CD8^-^ T cell (e.g. DN T, γδT cells, iNKT or MAIT) plays a role in Myc OFF-induced PDAC regression. The other cells that rapidly infiltrate PDAC tumours following Myc OFF are CD11c+, most commonly in mice these are cDC1 Dendritic cell cells but also mark age B cells and inflammatory macrophages. We used a *KM:Batf3^-/-^* mouse chimeric bone marrow transplant strategy to confirm that cDC1 cells do indeed play an essential role in the execution of the Myc-dependent pancreatic tumour regression programme. Accordingly, amidst their many roles^32^, dendritic cells are widely associated with debris clearance, remodelling and resolution after injury or infection through their release of resolvins, TGFβ, reparative cytokines and, especially, of IL-10^42,43^. IL-10 in turn acts as a central switch in the transition from inflammation to repair by promoting phenotypic switching in macrophages from pro-inflammatory to reparative, promoting efferocytosis and clearance of debris, limiting scarring and fibrosis, and supporting the rebuilding and remodelling of tissue through release of TGF-β, Amphiregulin (AREG), and VEGF^44^.

The inherent rapidity of the MycER*^T^*^2^ switchable system has allowed us to explore how de-activation of Myc in ongoing pancreatic adenocarcinomas engages an innate tissue morphogenic programme that instructs PDAC regression. In an accompanying manuscript (Kortlever *et al*., 2026) we also identify an analogous process in lung whereby inhibition of Myc rapidly engages a regression programme but is alerted by release of interleukin 33 (IL-33). In both tissues, de-activation of Myc is rapidly followed by release of an initiating alert signal around 2-6 hours later: GM-CSF in pancreatic adenocarcinomas and IL-33 in lung adenocarcinomas. The alert signal is received by sentinel cells — dendritic cells in pancreas and eosinophils in the lung — which then propagates a signalling cascade across the tumour that drives extensive remodelling, clearance of debris and removal of supernumerary cells. We have shown here that the rapidity of onset, complexity and reproducibility of tumour regression process has all the qualities of wound resolution — the final stage of the healing process where the integrity of a damaged tissue is restored and returned to its homeostatic state: this is not oncogene addiction, but redirection.

## METHODS

### Generation and maintenance of genetically engineered mice

C57BL/6J *LSL-Kras^G12D^*, *p48^cre/+^* and *Rosa26-lsl-MER^T^*^2^ (*KM*) mice have all been previously described^26,45–49^. Mice were maintained on a 12-hour light/dark cycle with continuous access to food and water and in compliance with protocols approved by the UK Home Office guidelines under project licenses to GIE at the Cambridge Research Institute (C4750/A12077) and at The Francis Crick Institute (C4750/A19013A). Deregulated Myc activity was induced in pancreatic epithelia of 11 to 13-weeks old *KM* mice by tamoxifen diet (Harlan Laboratories UK, TAM400 diet) as described in ^27^ and control mice were maintained on regular diet. C57BL/6J *Batf3-/-* mice where kindly donated by the Reis e Sousa C. laboratory (The Francis Crick Institute).

### Tissue preparation and histology

Mice were euthanized by cervical dislocation and cardiac perfused with PBS when required. Pancreata were removed, fixed 24 hours in 10% neutral-buffered formalin, stored in 70% ethanol and processed for paraffin embedding. Tissue sections (4 μm) were stained with haematoxylin and eosin (H&E) using standard reagents and protocols (Sigma-Aldrich, GHS232 and HT110216).

### Immunohistochemistry, immunofluorescence and RNAscope

For immunohistochemical (IHC) and immunofluorescence (IF) analysis, paraffin-embedded sections (4 μm) were de-paraffinized, rehydrated and either boiled in 10 mM citrate buffer (pH 6.0) or treated with 20 µg/ml Proteinase K to retrieve antigens, depending on the primary antibody used. Primary antibodies used: rabbit monoclonal anti-Cytokeratin 19 (Abcam, ab133496), rabbit monoclonal anti-CD3 (SP7) (ThermoFisher-Scientific, RM-9107-RQ), rabbit monoclonal anti-CD11c (D1V9Y) (Cell Signaling, 97585), goat polyclonal anti-MMR/CD206 (R&D Systems, AF2535), rat monoclonal anti-neutrophils (7/4) (Cedarlane, CL8993AP), goat polyclonal anti-NKp46/NCR1 (R&D systems, AF2225), rat monoclonal anti-CD45R/B220 (RA3-6B2) (ThermoFisher-Scientific, MA1-70098), rabbit monoclonal anti-Ki67 (SP6) (ThermoFisher-Scientific, MA5-14520), mouse monoclonal anti-Pan-Keratin (C11) (Cell Signaling, 4545), rabbit polyclonal anti-GM-CSF (Novus, NB600-632) and rabbit monoclonal anti-CC3 (5A1E) (Cell Signaling, 9664). Primary antibodies were incubated with sections overnight at 4°C. For IHC analysis, primary antibodies were detected using Vectastain Elite ABC HRP Kit (Vector Laboratories, Peroxidase Rabbit IgG PK-6101) and DAB substrate (Vector Laboratories, SK-4100); slides were then counterstained with haematoxylin solution (Sigma-Aldrich, GHS232) and mounted in DPX Mountant (Sigma-Aldrich, 06522). For IF analysis, primary antibodies were visualized using species-appropriate cross-adsorbed secondary antibody Alexa fluor 488 (ThermoFisher-Scientific A11008, A11006, A11001, A11055) or 555 conjugates (ThermoFisher-Scientific A21428, A21434, A21422, A21432); slides were counterstained with Hoechst (Sigma-Aldrich, B2883) and mounted in Prolong Gold Antifade Mountant (ThermoFisher-Scientific, P36934). TUNEL staining was performed using the Apoptag Fluorescein *in situ* Apoptosis Detection Kit (Millipore, S7110) according to manufacturer’s instructions. RNA-*in situ* hybridization (RNA-ISH) for mouse *Rasd2* (ACD, 416582) was amplified with RNAscope 2.5 HD Reagent Kit (Advanced Cell Diagnostics, 322300) and developed with the TSA Plus kit (PerkinElmer, NEL760001KT) according to manufacturer’s instructions. Images were collected with a Zeiss Axio Imager M2 microscope equipped with software ZEISS ZEN blue 3.2.

### Mouse Cytokine antibody Arrays

Pancreata were collected at indicated time points and snap-frozen in liquid nitrogen. Whole pancreas protein extracts were isolated and incubated with mouse inflammation/cytokine antibody arrays (R&D Systems, ARY028), according to manufacturer’s instructions. For IF analysis and quantification, arrays were visualized using IRDye 800CW secondary antibody (LI-COR, 926-32230). Signal intensity was acquired and analysed using LI-COR Odyssey CLX system.

### Pancreata isolation and quantitative real time-PCR

Pancreata were collected at indicated time points and snap-frozen in liquid nitrogen. Tissues were ground into powder in liquid nitrogen and total RNA was isolated using TRIzol Reagent (ThermoFisher-Scientific, 15596-018) following manufacturer’s instructions. cDNA was synthesised from 1 µg of RNA using the High-Capacity cDNA Reverse Transcription Kit with RNase Inhibitor (ThermoFisher-Scientific, 4374966) following manufacturer’s instructions. Quantitative reverse transcription PCR (qRT-PCR) reactions were performed on Applied Biosystems QuantStudio 5 Real-Time PCR System using Fast SYBR Green Master Mix (ThermoFisher-Scientific, 4385612) following manufacturer’s instructions. Primer used: mRasd2-F AACTGCGCCTACTTCGAGG, mRasd2-R GGTGAAAAGCATCGCCGTACT, mActin-F GACGATATCGCTGCGCTGGT, mActin-R CCACGATGGAGGGGAATA.

### Pancreata isolation, single cell RNA sequencing and analysis

Pancreata were isolated as described previously^50^, but with modifications. Briefly, mice were euthanized at indicated time points, hearts were flushed and pancreata collected in PBS containing 10% FBS, 1% Glucose, 0.1 mg/ml Soybean trypsin Inhibitor (Sigma-Aldrich, T6522) and Protease inhibitor cocktail tablet (Roche, 11873580001). Samples were weighed, mechanically dissociated and digested in 5 ml of Hanks’ balanced salt solution (HBSS) containing 750 U/ml collagenase I (ThermoFisher-Scientific, 17018029), 0.3 mg/ml DNase I (Sigma-Aldrich, DN25) and 0.1 mg/ml Soybean trypsin inhibitor for 45 min at 37°C on a shaker (200 rpm), followed by dissociation with a syringe and needle and filtration through a 70-μm strainer. Cells were stained using DRAQ5 (ThermoFisher-Scientific, 62252), Fixable Viability Dye eFluor 780 (eBioscience, 65-0865), and sorted as Live single cells on a FACS Aria instrument (BD Biosciences).

Library preparation of murine samples was performed according to instruction in the 10X Chromium single cell kit 3’ v3. Libraries were sequenced on a NovaSeq6000 NGS platform. Read processing was performed using the 10X Genomics workflow. Briefly, Cell Ranger Single-Cell Software Suite (6.1.1) was used for demultiplexing, barcode assignment and UMI quantification (http://software.10xgenomics.com/single-cell/overview/welcome). The reads were aligned to the pre-built mm10 reference genome provided by 10X Genomics with the *MYCER^T^*^2^ fusion gene sequence added. Raw single-cell sequencing data from individual samples were processed using the 10x Cell Ranger pipeline (10X Genomics) and analysed with the Seurat R-package (version 5.0.0)^51^ in R version 4.3.1. Cells were filtered to exclude cells with mitochondrial gene percentages higher than 25% and cells with less than 200 RNA features per cell. For sample integration, the R-package harmony was used^52^. In order to create the UMAP plots, the first 50 PCA dimensions were included, and cell-cycle regression was performed. The FindClusters function from the Seurat R-package was used to define clusters with the cluster parameter set to 0.5. The python package decoupler was used to assign cell type identities to clusters using the decoupler-py python package^53^ in conjunction with custom cell types and marker gene sets. Single-cell differential gene expression analyses were conducted with the glmGamPoi R-package (version 1.12.2)^54^. RNA-seq data have been deposited in the Gene Expression Omnibus database at NCBI (https://www.ncbi.nlm.nih.gov/geo/) under accession number GSE319823.

### Blocking antibody and recombinant protein treatments

For GM-CSF blocking studies, *KM* mice were *i.p.* injected with 250 µg of rat monoclonal anti-mouse GM-CSF antibody (MP1-22E9) (BioCell, BE0259) or equivalent amount of rat IgG2a isotype control (2A3) (BioCell, BE0089) every 2 days, starting one day prior to MycER deactivation. Mice were euthanized at the indicated time points and pancreata harvested for histological analysis.

For GM-CSF recombinant studies, *KM* mice were administered tamoxifen diet for 17 days. Commencing at 14 days mice were *i.p.* injected with 2 doses 24 hours apart of 3 µg of carrier free recombinant mouse GM-CSF (R&D systems, 415-ML-CF) reconstituted in sterile PBS. Control mice were injected with an equivalent amount of PBS. Mice were euthanized 3 days later and pancreata harvested for histological analysis.

### Bone marrow chimeras

cDC1-specific KO mice with inducible epithelial tumours were generated as previously described^34^. Briefly, 6 to 8-week-old CD45.2+;*KM* mice were lethally irradiated with two doses of 6.6 Gy (3-4 hours rest in between exposures). Mice were injected the next day *i.v.* with bone marrow cells from CD45.1+;*Batf3-/-* or CD45.1+;*WT* mice. Chimerism was quantitated 5 weeks later from splenocytes using flow cytometry protocol described below. 14 weeks old chimeric mice were subjected to tamoxifen diet treatment and withdrawal as described above and in the Results section. Samples were collected at the indicated time points.

### Flow cytometry

For pancreatic immune cell detection: single cell suspension was performed as describe above and Fc receptors were blocked using anti-mouse CD16/CD32 (Biolegend). Dead cells were excluded with the fixable viability dye eFluor 455UV (eBioscience) and the following fluorescent label-conjugated antibodies were used: CD45 (30-F11, Biolegend, BV510), SiglecF (1RNM44N, eBioscience, SB600), XCR1 (ZET, Biolegend, BV650), CD11b (M1/70, eBioscience, BV786), MHCII (CI2G9, BD Biosciences, BUV395), CD172a (P84, Biolegend, Alexa Fluor 488), Ly6G (1A8-Ly6g, eBioscience, PE-eFluor 610), Ly6C (HK1.4, eBioscience, PE-Cy7), CD11c (N418, eBioscience, Alexa Fluor 700), F4/80 (BM8, eBioscience, APC-eFluor 780), and lineage cocktail which contained CD5 (53-7.3), CD19 (1D3), NK1.1 (PK136), CD3e (145-2C11), B220 (RA3-6B2) (eBioscience, eFluor 450).

For splenic immune cell detection: single cell suspension and staining was performed as previously described^55,56^. Briefly, splenocytes were digested with 400 U/mL collagenase IV (Worthington, LS004209) and 0.4 mg/mL deoxyribonuclease I (Sigma-Aldrich, 11284932001) in RPMI 1640 for 15 minutes. Digested tissues were strained through a 70 µm strainer and washed and resuspended with FACS buffer (PBS + 3% FBS, 2 mM EDTA). Single cell suspensions were stained in PBS with LiveDead Fixable dye (Thermo Fisher Scientific) according to manufacturer’s instructions, Fc block (anti-CD16/32) (BD Pharmingen, 2.4G2) and subsequently stained with the following fluorescent label-conjugated antibodies: CD45R/B220 (Biolegend, RA3-6B2), I-A/E (Thermo Fisher Scientific, M5/114.15.2), CD11c (Biolegend, N418), XCR1 (Biolegend, ZET), CD172a/Sirpα (Biolegend, P84). All data were acquired on a Fortessa or Symphony instrument (BD Biosciences) and analysed using FlowJo X (Tree Star).

### Images processing, quantification and statistical analysis

For quantification of tumour burden (H&E) and Krt19 IHC expression the stained sections were scanned with an Aperio AT2 microscope (Leica Biosystems) at 20X magnification (resolution 0.5 μm per pixel) and analysed with Aperio Software v12.4.0.5043. IHC and IF stained slides were analysed and quantified by randomly selecting a minimum of 5 pictures per sample and counting the positively identified cells per tumour field of view (FoV) using Fiji ImageJ 1.52i open-source software. Statistical significance was determined on the median per condition/group values by unpaired non-parametric Multiple Mann-Whitney U test or unpaired Student’s t-test with Welch’s correction using Prism GraphPad software versions 9 and 10. Quantifications in scatter column bar and dot-plot graphs show median with interquartile range (+/- upper and lower quartiles) and display p-values to four decimal places. Quantification of qPCR shown as XvY graph include a non-linear regression (fit curve), half-life=3.1.

## Data availability

Raw single cell sequencing data generated in this study were deposited in the NCBI Gene Expression Omnibus (accession number GSE319823). All other relevant data supporting the key findings of this study are available within the Article and its Extended Data Figures or from the corresponding author upon reasonable request.

## AUTHOR CONTRIBUTIONS

TC and GIE conceived the project. TC, RMK and GIE designed experiments. TC performed all experiments. RMK, NMS, MDB, JS, JP, CML, JRPM, AP and TYFH assisted with some experiments. EA and BS assisted with the bioinformatic analysis. TC, RMK and GIE wrote the manuscript. GIE supervised the study. All authors discussed results and revised the manuscript.

## ACKNOWLEDGEMENTS

We thank past and present members of the Evan laboratory for invaluable discussion and advice. We also thank Irina Pshenichnaya (Cambridge Stem Cell Institute) for the RNAscope staining and Caetano Reis e Sousa for helpful discussion and suggestions.

This study was supported by programme grants to GIE (Cancer Research UK C4750/A12077, C4750/A19013A and the SU2C/CRUK/Lustgarten Foundation Transatlantic Pancreatic Cancer Dream Team C4750/A22585) and H2020 Marie Skłodowska-Curie Actions Individual Fellowships awarded to JS and MDB (84050 and 837951, respectively).

## ADDITIONAL INFORMATION

Correspondence and request for material can be sent to T. Campos and G.I. Evan. Reprints and permissions information is available at www.nature/com/reprints.

## FIGURE LEGENDS

**Extended Data Figure 1. PDAC regression after Myc OFF associated with cDC1 influx**

**a.** Left: Cell death (TUNEL) immunofluorescence and cell proliferation (Ki67) immunohistochemistry quantification of pancreata from *KM* mice treated with normal food (-), tamoxifen food (Myc ON) for 2 weeks (+) or tamoxifen food for 2 weeks followed by normal food (then Myc OFF) for the indicated time points. N = 5-8 mice per group. Shown are means and standard deviations per group. Statistical analysis was performed using multiple unpaired *t*-tests with Welch’s correction. Right: representative cell death (TUNEL; top) and proliferation (Ki67; bottom) immunohistochemistry of pancreata from *KM* mice treated with normal food (Myc never ON), tamoxifen food for 2 weeks (Myc ON) or tamoxifen food for 2 weeks followed by normal food for 5 days (Myc OFF 5d). Scale bar across the row = 100 μm.

**b.** Representative CD11c immunofluorescence staining of pancreata from *KM* mice treated with normal food (Myc never ON), tamoxifen food for 2 weeks (Myc ON) or tamoxifen food for 2 weeks followed by normal food for 5 days (Myc OFF 5d). Scale bar across the row = 50 μm.

**c.** Left: H&E staining quantification of pancreata from *M*, *K* and *KM* mice treated with tamoxifen food (TAM) for 2 weeks (+), normal food (-) or tamoxifen food for 2 weeks followed by normal food (then no TAM) for the indicated time points. N = 5-8 mice per group. Shown are medians with interquartile ranges per group. Statistical analysis was performed using multiple Mann-Whitney tests. Right: representative haematoxylin and eosin (H&E) staining of pancreata from *KM* mice treated with normal food (Myc never ON), tamoxifen food for 2 weeks (Myc ON) or tamoxifen food for 2 weeks followed by normal food for 5 days (Myc OFF 5d). Scale bar across the top row = 2 mm. Bottom row shows higher magnification of the corresponding insets in top panels, scale bar across the row = 100 μm. L= lesion area.

**Extended Data Figure 2. *Rasd2* is a biomarker for Myc activity**

**a.** RNA-sequencing in pancreatic tumour cell line from *KM* mouse. Heatmap shows all downregulated genes with FDR<0.01 upon Myc de-activation at all 3 indicated time points. Genes were ranked based on their expression level in the 6 hours Myc OFF versus Myc ON comparison. N = 3 samples per time point. Statistical analysis was performed using multiple testing via the FDR (Benjamini-Hochberg approach).

**b.** Genome browser tracks showing Myc-ChIP signal in the *Rasd2* promoter (shaded area) in *KM* PDAC cell line treated with 4-OHT (Myc ON), or 4-OHT followed by vehicle for 1 day (Myc OFF 1d) or 2 days (Myc OFF 2d).

**c.** Representative RNA *in situ* hybridisation of *Rasd2* (red) in pancreata from *KM* mice treated with tamoxifen food for 2 weeks (Myc ON) or tamoxifen food for 2 weeks followed by normal food (Myc OFF) for the indicated time points. Scale bar across the top row = 100 μm. Bottom row shows higher magnification of the corresponding insets in top panels, scale bar across the row = 20 μm

**d.** qRT-PCR analysis of *Rasd2* expression in mRNA isolated from *KM* mice treated with tamoxifen food for 2 weeks (Myc OFF 0 hours) or tamoxifen food for 2 weeks and then normal food for the indicated time points. *β-Actin* was used as internal amplification control. Dotted line indicates *Rasd2* half-life (3.1 hours). N = 3 samples per time point. Statistical analysis of the XvY graph was performed using a non-linear regression (fit curve).

**Extended Data Figure 3. Single cell and cytokine analysis of PDAC cells show rapid and MycER*^T2^*+ ductal-specific induction of GM-CSF after Myc OFF**

**a.** Dot plot of marker genes used to identify subpopulations in the single-cell RNA-sequence data from pancreata of *KM* mice treated with tamoxifen food for 2 weeks or tamoxifen food for 2 weeks followed by normal food for 6 hours.

**b.** UMAP representation of scRNA-sequence analysis of pancreata from *KM* mice treated with tamoxifen food for 2 weeks and tamoxifen food for 2 weeks followed by normal food for 6 hours (see Fig. 2b). *MycER^T^*^2^ expression is highlighted in red.

**c.** Heatmap representation of ductal subpopulations showing cluster-specific differentially upregulated genes. Cluster identity (C0-C6) relates to the subclustering UMAP in Fig 2c.

**d.** Representative immunofluorescence staining of Pan-Keratin (PanK; green) and granulocyte-macrophage colony-stimulating factor (GM-CSF; red) in pancreata harvested from *KM* mice treated with tamoxifen food for 2 weeks (Myc ON) or tamoxifen food for 2 weeks followed by normal food for 6h (Myc OFF 6h). Scale bar across the row = 100 μm.

**e.** Quantification of GM-CSF measured using Cytokine array analysis of pancreata from *KM* mice treated with normal food (-), tamoxifen food (Myc ON) for 2 weeks (+) or tamoxifen food for 2 weeks followed by normal food for the indicated time points (then Myc OFF). N = 4-6 mice per group. Shown is median with interquartile range per timepoint.

**Extended Data Figure 4. Immune microenvironment responses of PDAC in first 10 days after Myc OFF**

**a-e.** Quantification of immunostainings for **a**, T cells (CD3), **b**, B cells (B220), **c**, Neutrophils (Ly6b), **d**, NK cells (NKp46) and **e**, Macrophages (CD206) in pancreata harvested from *KM* mice treated with tamoxifen food (Myc ON) for 2 weeks (1^st^ column, data taken from Fig. 1c) or tamoxifen food for 2 weeks followed by normal food for 3 or 10 days while treated with IgG control (2^nd^ and 4^th^ column) or granulocyte-macrophage colony-stimulating factor (αGM-CSF) antibody (3^rd^ and 5^th^ column). FoV = field of view. N = 5-8 mice per group. Shown are medians with interquartile ranges per condition/timepoint. Statistical analysis was performed using multiple unpaired *t*-tests with Welch’s correction.

**Extended Data Figure 5. Immune microenvironment responses of PDAC component cells in first 3 days after rGM-CSF therapeutic treatment**

**a-e.** Quantification of immunostainings for **a**, T cells (CD3), **b**, B cells (B220), **c**, Neutrophils (Ly6b), **d**, NK cells (NKp46) and **e**, Macrophages (CD206) in pancreata harvested from *KM* mice treated with tamoxifen food for 17 days (1^st^ column) or tamoxifen food for 17 days and treated with rGM-CSF (2^nd^ column). N = 5-7 mice per group. FoV = field of view. Shown are medians with interquartile ranges per condition/timepoint. Statistical analysis was performed using multiple unpaired *t*-tests with Welch’s correction.

**Extended Data Figure 6. Analysis of cDC1 cells in chimeric mice and tumour cell death in *KM:WT* mice.**

**a.** Fluorescence-activated cell sorting (FACS) analysis of pancreata from *KM* mice treated with tamoxifen food for 2 weeks (1^st^ column) or tamoxifen food for 2 weeks followed by normal food for 3 days (2^nd^ column). Analysis of cDC1 cells as percentage of total cDC cells. N = 3-4 mice per group.

**b.** Fluorescence-activated cell sorting (FACS) analysis of spleens from chimeric *KM:WT* or *KM:Batf3^-/-^* mice treated with tamoxifen food for 2 weeks (1^st^ column) or tamoxifen food for 2 weeks followed by normal food for 3 days (2^nd^ column). Analysis of indicated cells as percentage of live CD45+ cells. N = 3-4 mice per group.

**c.** Staining quantification of pancreata from chimeric *KM:WT* mice treated with tamoxifen food for 2 weeks (1^st^ column) or tamoxifen food for 2 weeks followed by normal food for 3 days (2^nd^ column). Tissue sections stained for cell death (TUNEL). N = 3-4 mice per group.

**a, b, c:** Shown are median with interquartile range per condition/timepoint. Statistical analysis was performed using multiple unpaired *t*-tests with Welch’s correction.

**Extended Data Table 1.**
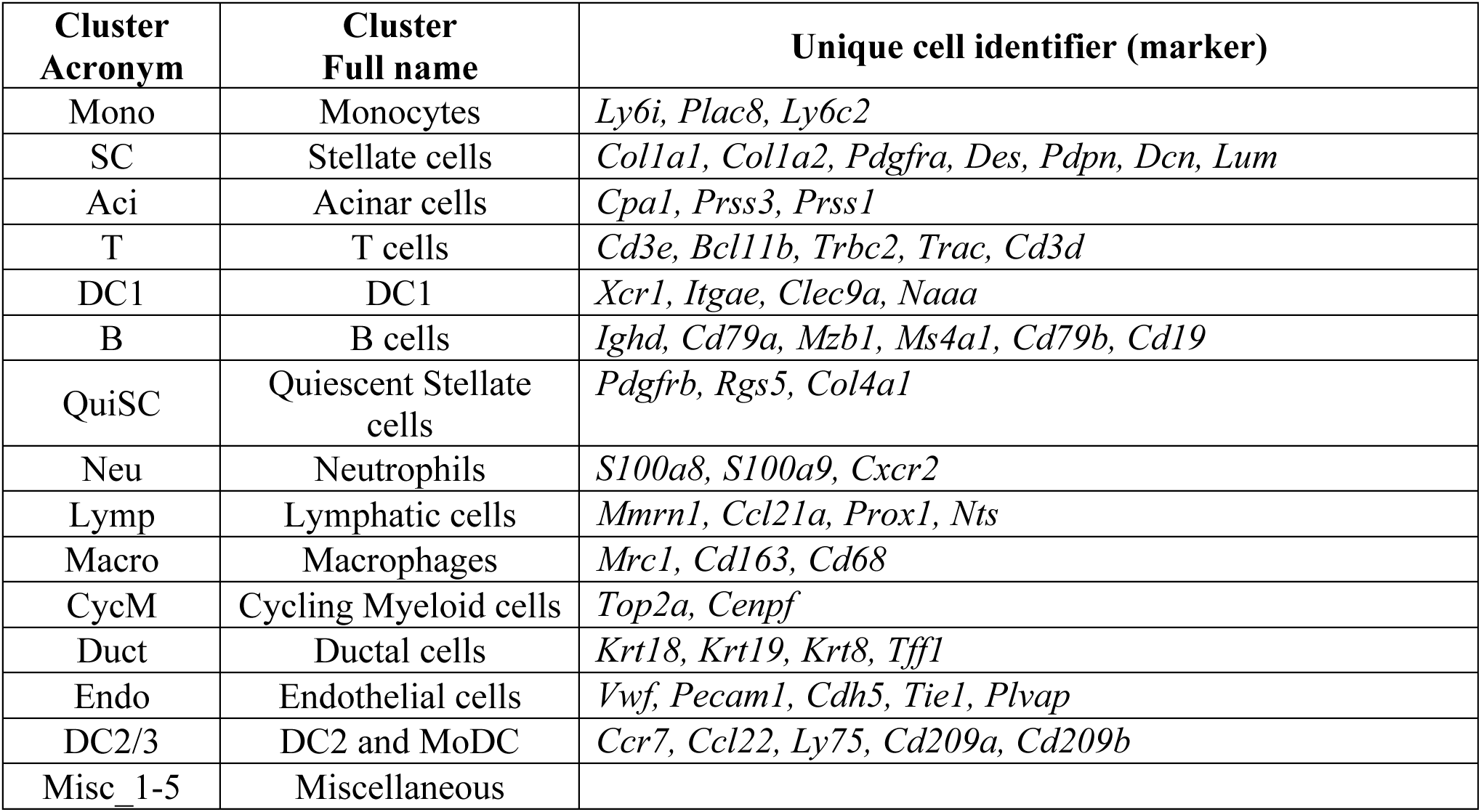

