## Supplemental files for "Myc inhibition triggers GM-CSF-driven regression of pancreatic tumours"

### Extended Data Figure 1

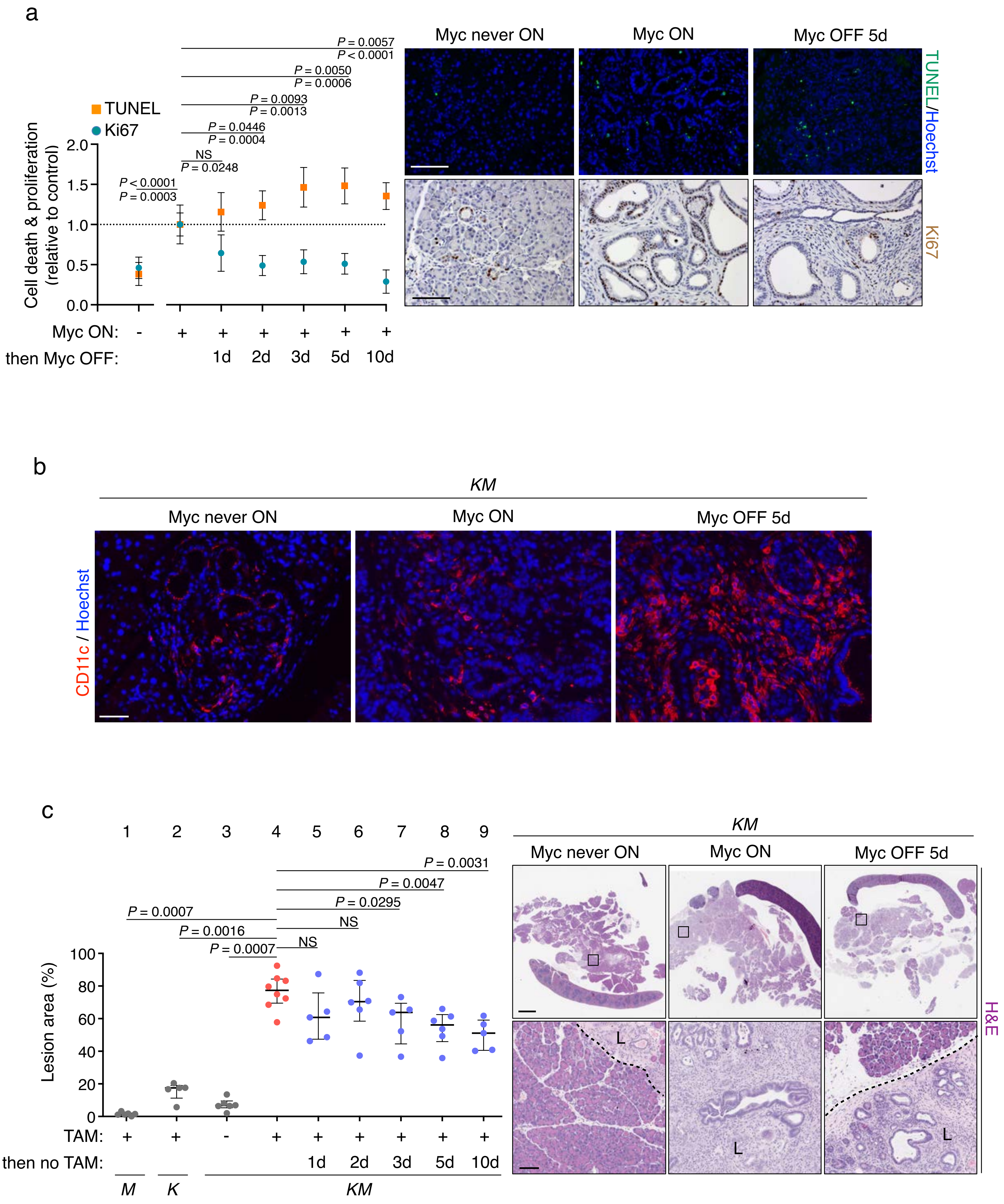

### Extended Data Figure 2

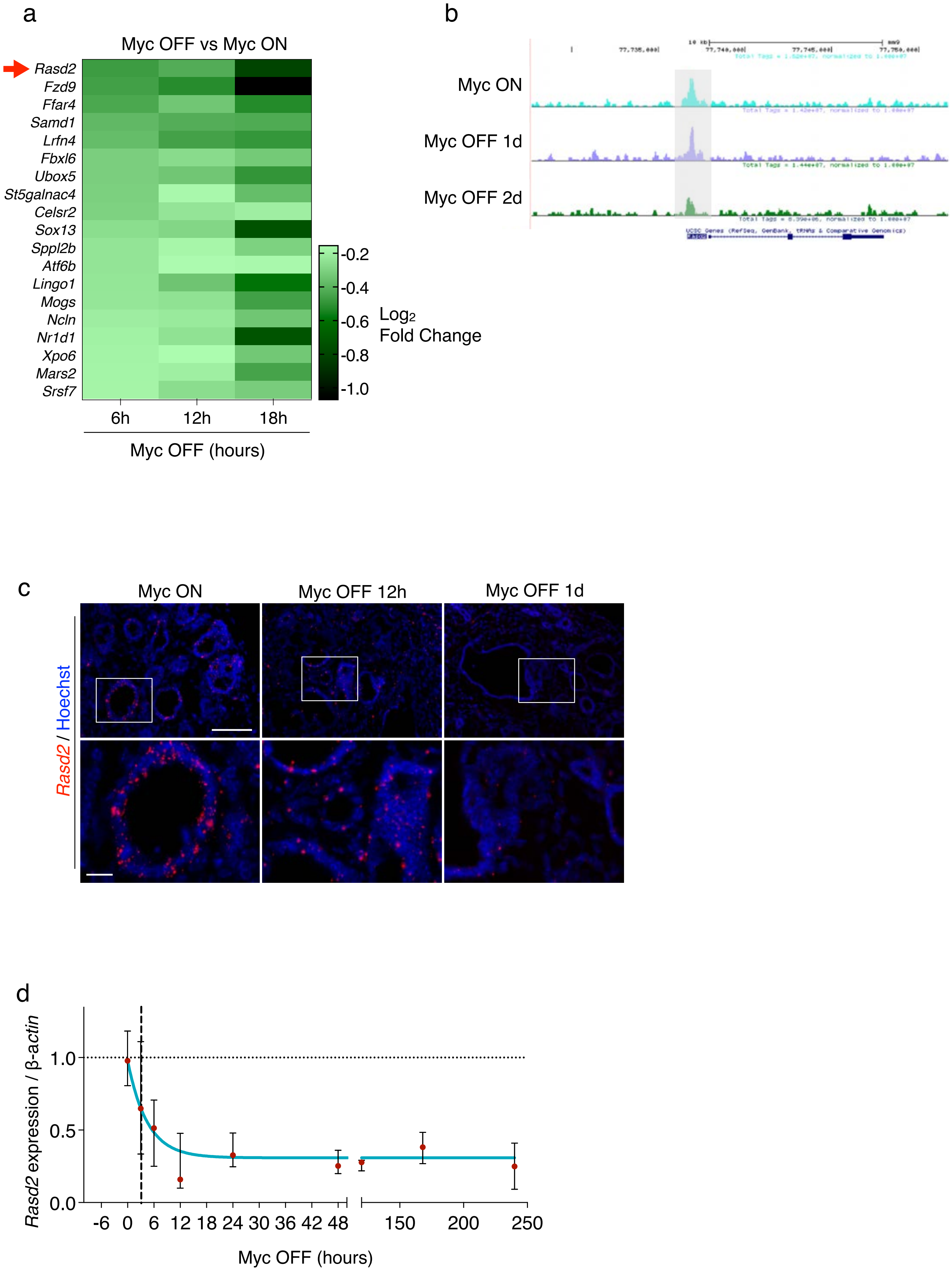

### Extended Data Figure 3

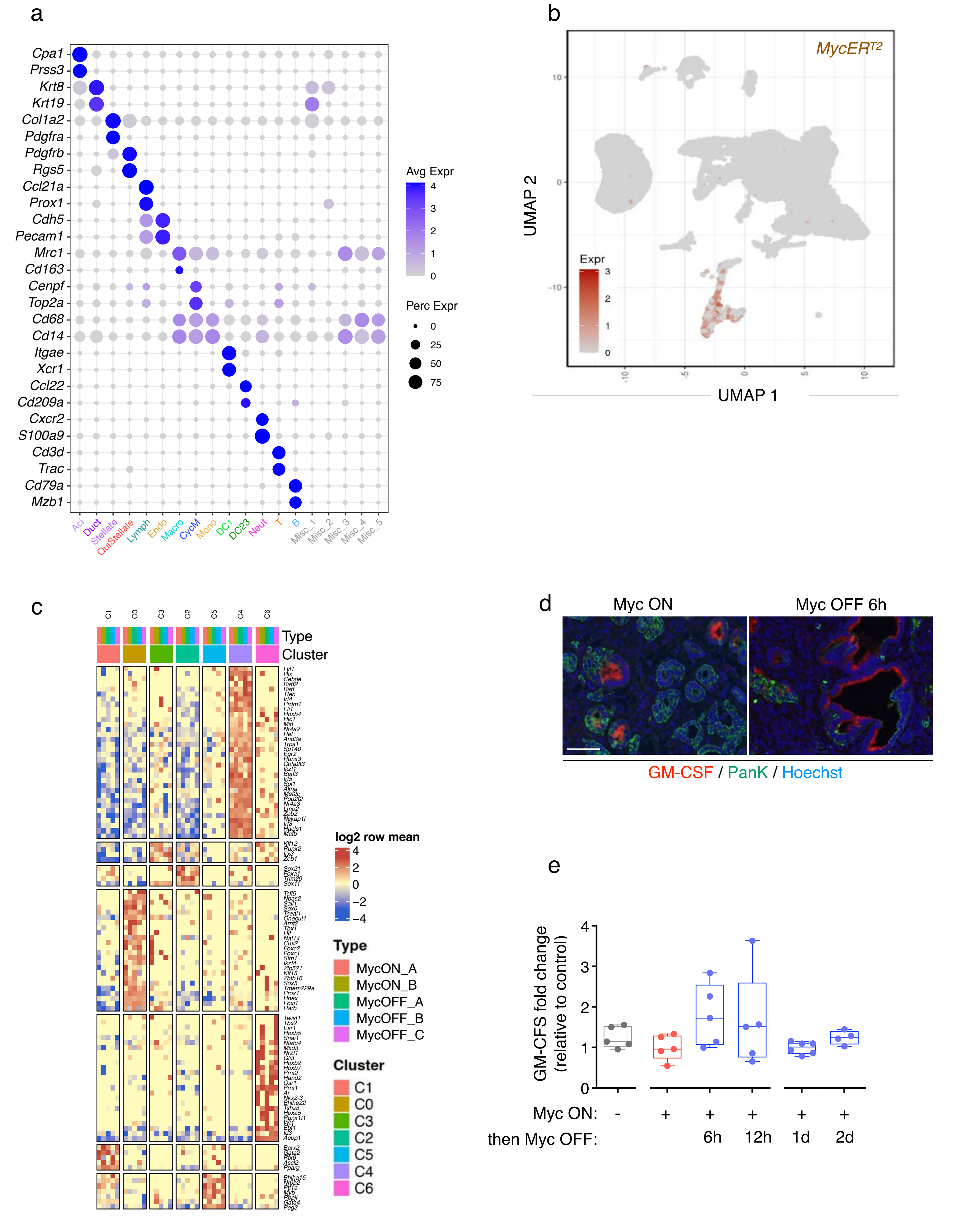

### Extended Data Figure 4

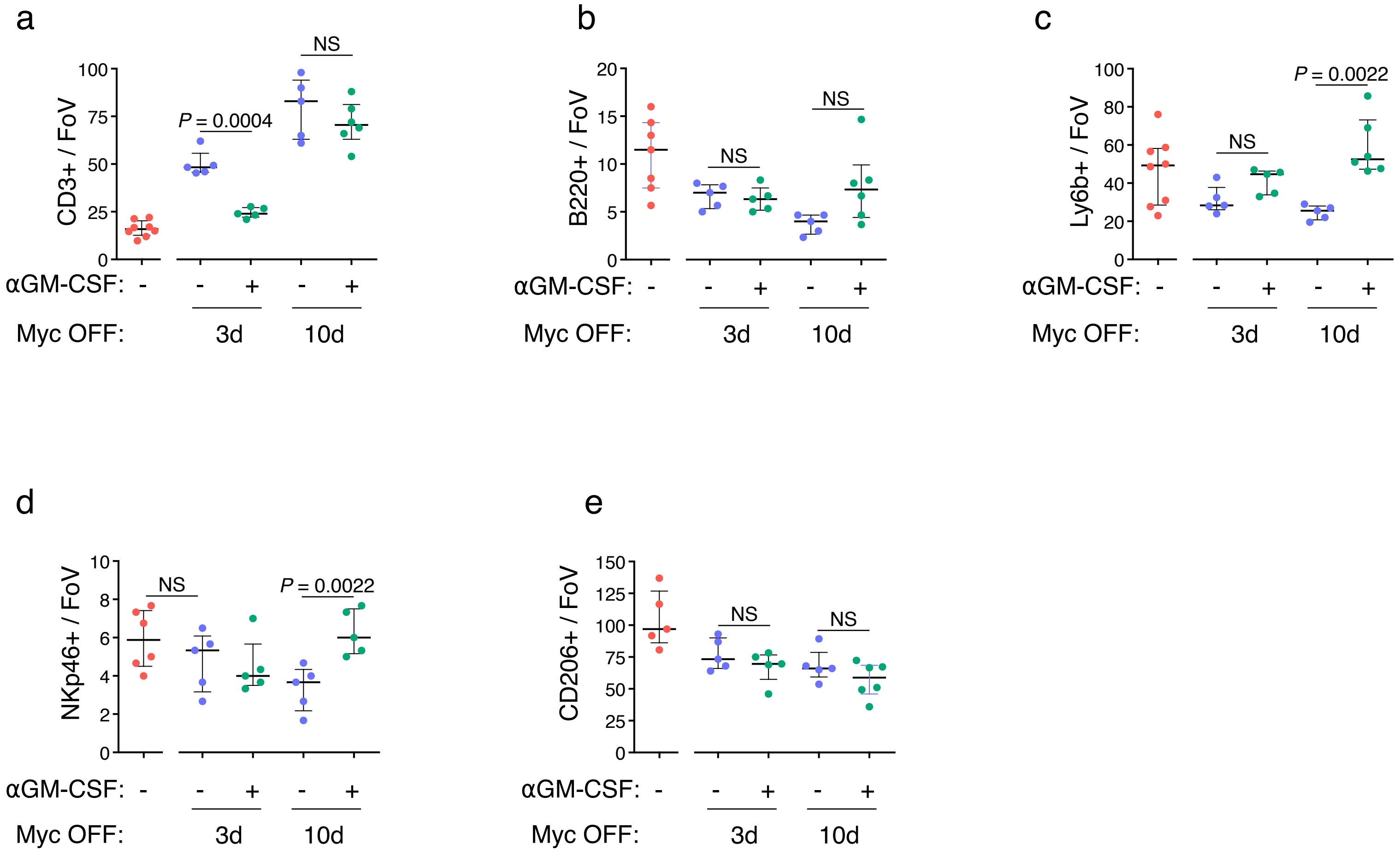

Extended Data Figure 5

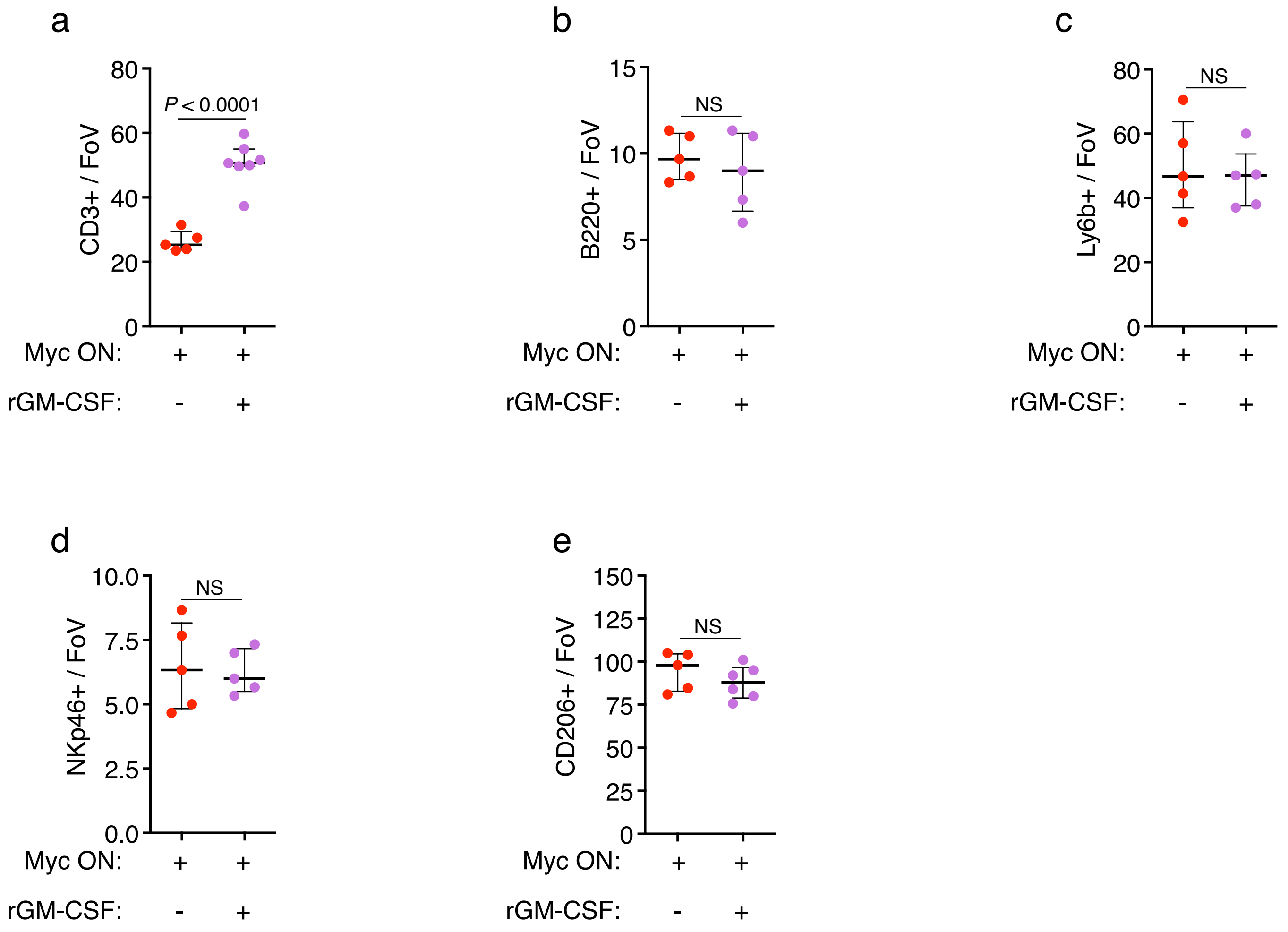

### Extended Data Figure 6

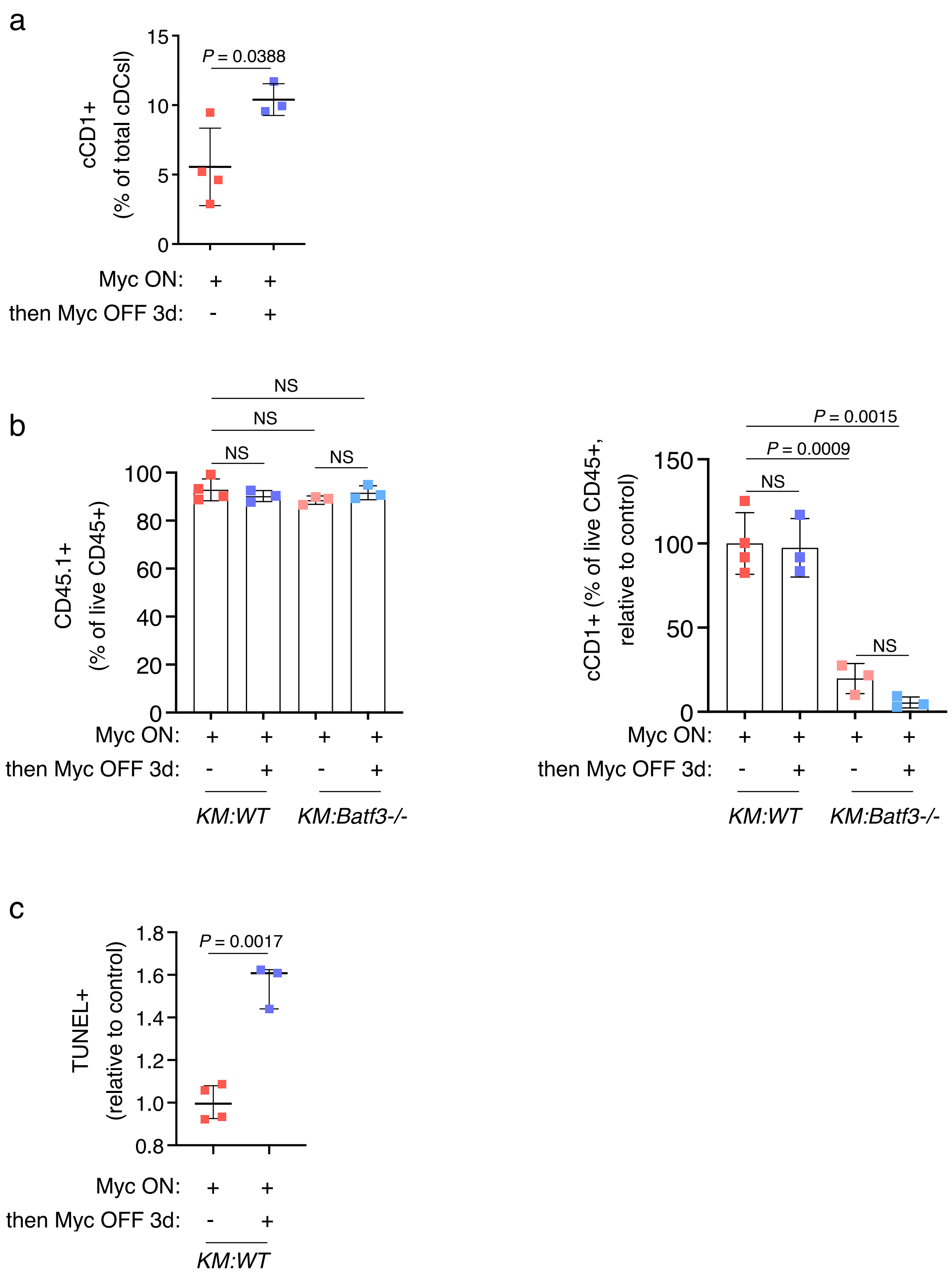
